# Identification and Characterization of Metastasis-initiating cells

**DOI:** 10.64898/2026.02.03.703643

**Authors:** Sijia Wu, Jiangpeng Wei, Xin Liu, Jiajin Zhang, Jianguo Wen, Liyu Huang, Xiaobo Zhou

## Abstract

Metastasis, the primary cause of cancer-related mortality, is a dynamic and complex process driven by a subset of cells known as metastasis-initiating cells (MICs). Accurate identification of MICs is therefore critical for metastasis diagnosis and therapeutic decision-making. However, current approaches rely either on mouse tracing experiments, which are difficult to translate to human systems, or on indirect strategies such as stemness, trajectory, pathway, and biomarker analyses that often yield inconsistent results. To address these limitations, we propose scMIC, a computational framework designed to explicitly and reliably identify MICs from single-cell data (available at https://github.com/swu13/scMIC). scMIC integrates an embedding-based representation, unbalanced optimal transport, and a top-k selection strategy to robustly capture metastasis-initiating potential. The framework was validated and applied across multiple cancer types, species, and multi-omics datasets. Our results demonstrate the reliability of scMIC for MIC identification, its potential clinical utility in metastasis prognosis, and its effectiveness in discovering metastasis-related gene programs and molecular biomarkers. Elucidating the mechanisms of metastasis initiation not only advances our understanding of metastatic progression but also enables the development of therapeutic strategies that target the more aggressive MIC population rather than non-MICs, thereby avoiding unintended increases in metastatic risk. Collectively, scMIC provides a powerful tool for cancer metastasis research and drug discovery.

## 1. Introduction

Metastasis, the ultimate manifestation of cancer, accounts for the majority of cancer-related mortality (1). It arises from a complex and often inefficient series of biological events collectively known as the metastatic cascade (2). During this process, only a small subset of tumor cells is capable of adapting to the hostile and dynamic microenvironment (3). These cells, termed metastasis-initiating cells (MICs), undergo extensive intrinsic and dynamic reprogramming (4). The abundance and functional capacity of MICs within a primary tumor can serve as a predictive indicator of metastatic potential and inform treatment strategies. Therefore, accurately identifying MICs within primary tumors and characterizing their intrinsic gene programs is critical for understanding metastatic progression.

Researchers have made substantial efforts to identify and characterize MICs. One approach involves establishing metastatic mouse models with multi-timepoint tracking (5), which can accurately identify MICs. However, this method is difficult to implement in humans. Despite its limitations, the high accuracy of the metastatic mouse model makes it an ideal gold standard for developing a computational framework for MIC identification. Another approach leverages clonal information captured by an inducible CRISPR-Cas9-based lineage recorder (6). This high-resolution lineage tracing can reveal aggressive cell states, providing valuable validation for computational predictions. Additionally, researchers have employed complementary strategies such as pseudotime inference (7), stemness scoring (8), pathway enrichment analysis (9), candidate MIC biomarkers (10) to gain insights into metastatic potential. Given the hybrid characteristics of MICs (6), the distinction between tumor-initiating cells and MICs (11,12), as well as the inconsistencies and limited availability of well-established MIC biomarkers and pathways, there is a clear need for a robust computational framework to systematically study MICs. Such a framework can integrate diverse data sources, leverage wide features, and overcome the limitations of experimental approaches, enabling a more comprehensive understanding of metastatic initiation.

In this study, we propose an optimal transport-based framework to identify MICs from primary tumor cells (Figure 1A). This framework is grounded in the concept that metastatic tumor cells originate from MICs, highlighting their intrinsic similarity, and that MICs tend to undergo minimal changes when establishing specific metastases (13). Optimal transport theory has been widely applied in computational biology (14-16), and here it provides a natural approach to determine the most efficient mapping of MIC distributions to distant metastatic tumor cells, given a defined cost function (17). We apply this framework to multiple datasets to demonstrate its effectiveness (Figure 1A), assess the clinical relevance of MICs (Figure 1B), delineate the MIC gene program (Figure 1C), and identify candidate MIC biomarkers across multiple scales (Figure 1D). These aspects will be discussed in detail, one by one, in the following sections.

**Figure 1.**
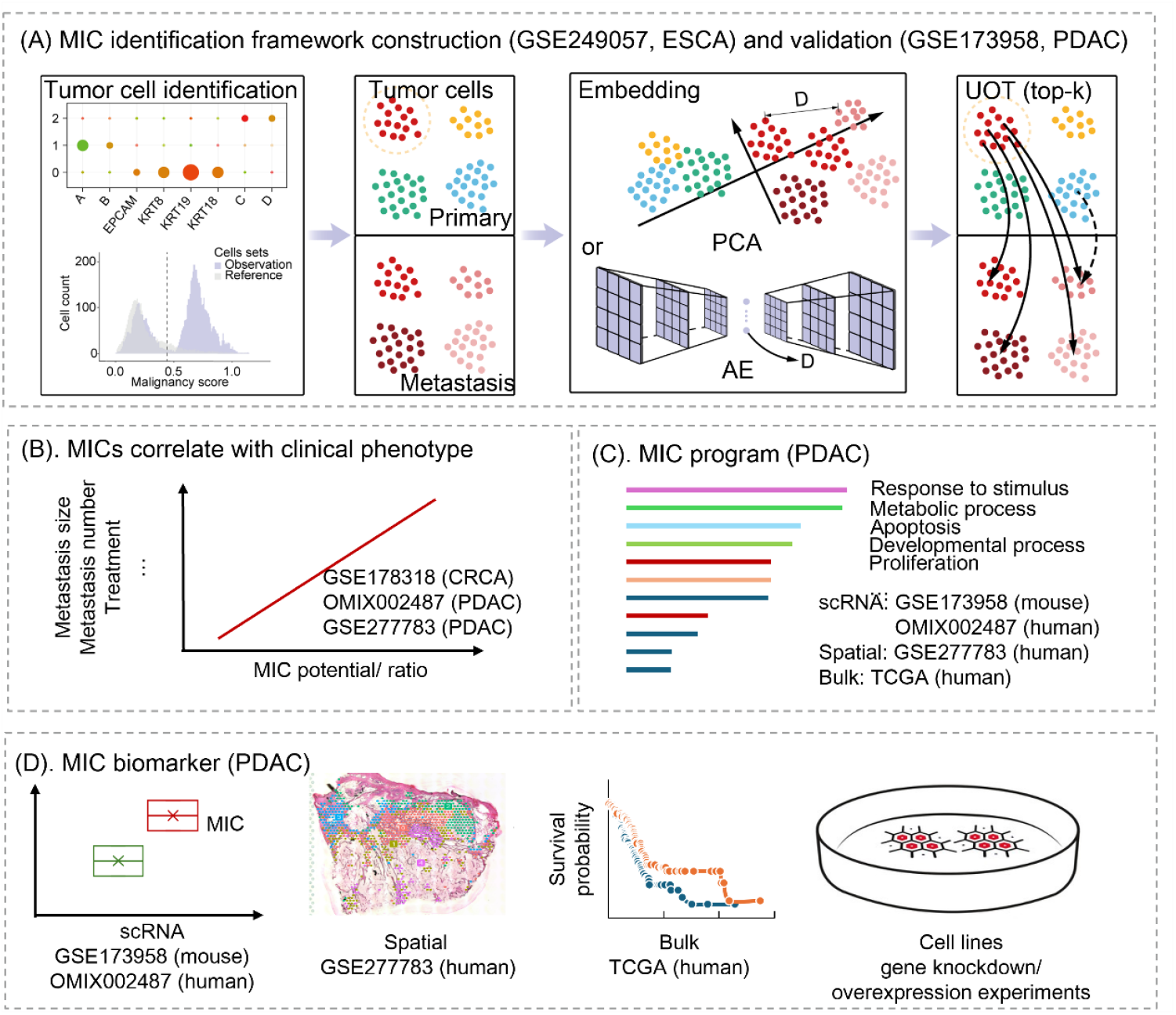
Study overview. (A) The scMIC framework for identifying metastasis-initiating cells (MICs) in esophageal squamous cell carcinoma (ESCA; GSE249057) with validation in pancreatic ductal adenocarcinoma (PDAC; GSE173958). (B) Associations between MICs and clinical phenotypes in colorectal cancer (CRCA, GSE178318) and PDAC (OMIX002487 and GSE277783). (C) Definition of the MIC transcriptional program across single-cell, spatial, and bulk transcriptomic data using PDAC as an example. (D) Multi-scale identification of MIC-specific biomarkers in PDAC. PCA, Principal Components Analysis; AE, AutoEncoder; UOT, Unbalanced Optimal Transport.

## 2. Results

### 2.1 Computational framework for MIC discovery

To construct a computational framework for MIC discovery, we first analyzed metastatic mouse models with multi-timepoint tracking data (GSE249057). The dynamic changes of lung metastatic cells across four timepoints (6 h, 48 h, 2 months, and 4 months) revealed a distinct cell population (Figure 2A). Specifically, cells in Cluster 1 decreased during the early metastatic stage and gradually rebounded at later stages, a trend consistent with the number of viable tumor cells measured by flow cytometry. Furthermore, Cluster 1 cells were markedly enriched in the subpopulations observed at 4 months with visible metastases in resected lungs. Taken together, these results indicated that Cluster 1 represented the MIC population. Notably, Cluster 1 cells exhibited dynamic epithelial– mesenchymal transition (EMT) activity, characterized by increased EMT at the early metastatic stage followed by decreased EMT at later stages (Figure 2B).

**Figure 2.**
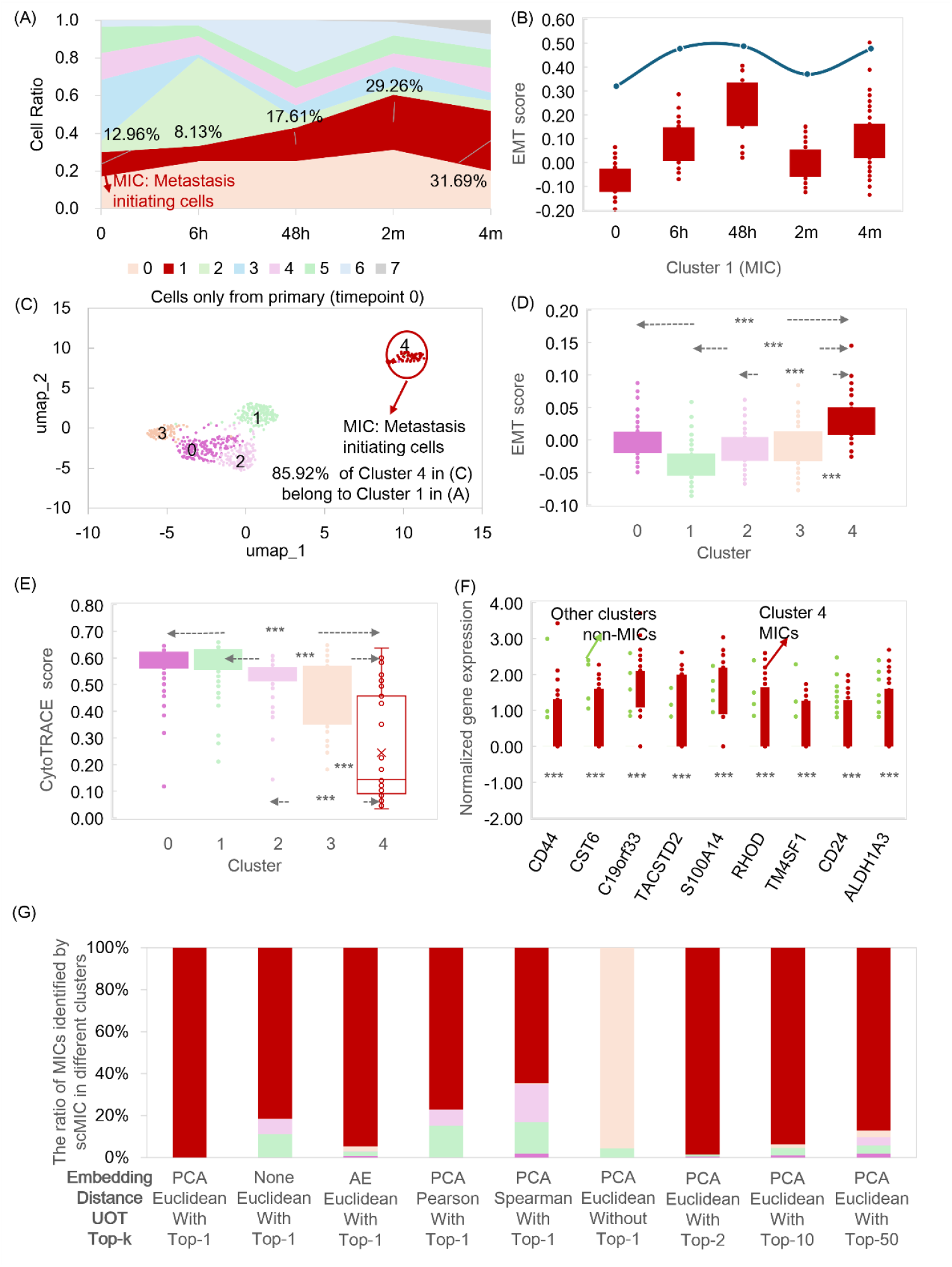
Training of MIC framework. (A) Multi-timepoint tracking of mouse tumors (GSE249057) identified cluster 1 as the MIC cluster (red). Numbers indicate the proportion of MICs at each timepoint. (B) Temporal dynamics of epithelial–mesenchymal transition (EMT) states within the MIC cluster during metastatic progression. (C) Re-clustering of primary tumor cells (timepoint 0) identified cluster 4 as the MIC cluster, consistent with multi-timepoint tracking results. (D) Comparison of EMT status between MIC and non-MIC clusters in primary tumors. (E) CytoTRACE scores of MIC versus non-MIC clusters in primary tumor cells. (F) Enrichment of metastasis-associated markers in the MIC cluster relative to non-MIC clusters in primary tumors. (G) Performance of the scMIC framework across embedding methods, distance metrics, UOT strategies, and Top-k selection schemes. Higher enrichment of MICs identified by scMIC in cluster 4 (red) indicates improved identification accuracy. ***, P < 0.001.

The primary tumor serves as the critical source of MICs. To investigate this, we clustered only parental tumor cells (0 h) and, by integrating the dynamic results across multiple timepoints, identified cluster 4 as the MIC population within the primary tumor (Figure 2C). Compared with other tumor cell populations, cluster 4 MICs exhibited a mesenchymal phenotype (Figure 2D), distinct stemness characteristics (Figure 2E), and elevated expression of metastasis-initiating signature genes previously reported in the literature (5) (Figure 2F). Interestingly, cluster 4 displayed a significantly lower CytoTRACE score, suggesting a more differentiated state, which contrasted with the conventional notion that MICs possessed higher stemness. Nevertheless, cluster 4 showed strong expression of CD44, a well-established stemness-associated marker. And CD44^high^ mice retained more viable tumor cells during metastatic progression than CD44^low^ counterparts. Taken together, these findings supported the robustness of cluster 4 as the MIC population in the primary tumor and underscored the need for a computational framework to systematically identify MICs.

Using parental tumor cells and lung metastatic cells at 4 months, we developed an optimal transport-based approach, scMIC (Figure 1A), to identify MICs in the primary tumor. Although the input scRNA-seq data here contained only tumor cells, to enhance its robustness and applicability, the first step of scMIC was to identify tumor cells based on marker expression, copy number variation (CNV), and other malignancy score (18). Next, we selected either significantly differentially expressed genes across tumor subclusters or the highly variable genes, and projected these features onto top principal components (PCs) via PC analysis (PCA) or onto latent representations via an AutoEncoder to reduce dimensionality. Cell-cell distances were then calculated using Euclidean, which served as the defined cost function. Unbalanced Optimal Transport (UOT) was subsequently applied to derive the optimal transport plan from primary tumor cells to metastatic tumor cells. To reflect the unique origin of metastatic cells, a top-1 filtering scheme was implemented, in which each metastatic cell was assigned only to the primary tumor cell with the highest transport probability, while all other associations were set to zero. Finally, each primary tumor cell received a score ranging from 0 to the total number of metastatic cells. After normalization by the number of metastatic cells or the maximum value, or alternatively without normalization, this score represented the metastatic potential of each primary tumor cell, with higher values indicating a greater likelihood of being an MIC. Figure 2G and Figure S1 illustrated the impact of various parameters on MIC identification, highlighting the critical roles of UOT and the top-1 filtering scheme. Consistently, the MICs identified by scMIC were confined to cluster 4 (Figure 2G), further supporting the accuracy and robustness of this framework in MIC identification.

### 2.2 MICs overlap with aggressive states

To further validate the robustness of scMIC in MIC identification, an independent mouse scRNA-seq dataset with clonal information obtained through an inducible CRISPR-Cas9-based lineage recorder (GSE173958) was analyzed (Figure 3A, Figure S2A). Lineage tracing identified clone 1 in mouse 1 and clone 2 in mouse 2 as the most aggressive clones. Consistently, compared with non-MICs and the overall tumor population, MICs were significantly enriched within these aggressive clones (Figure 3B, Figure S2B).

**Figure 3.**
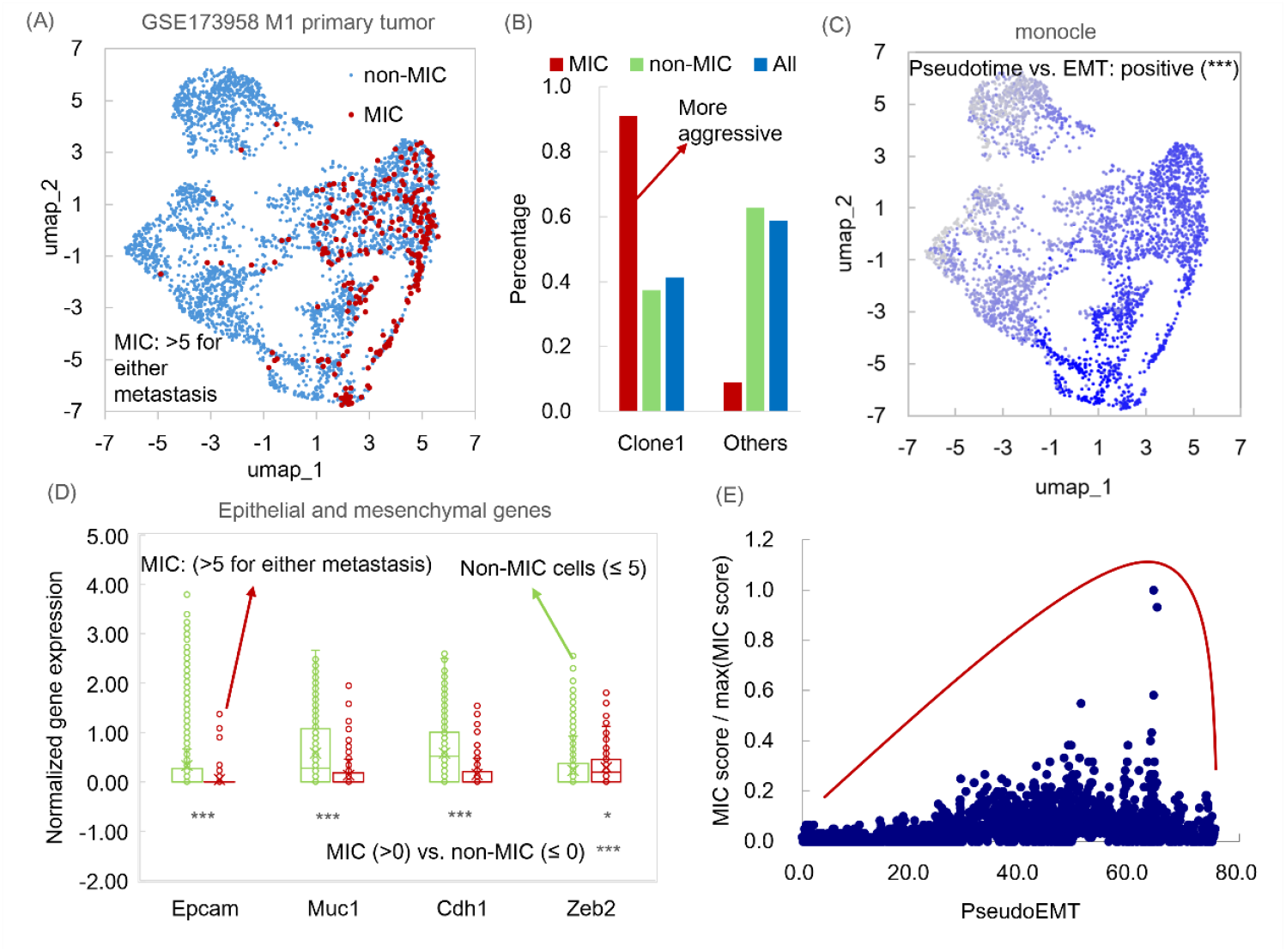
MICs associate with an aggressive clone and a hybrid EMT state in a lineage-traced M1 mouse model from the GSE173958 dataset. (A) UMAP visualization of primary tumor cells with MICs identified by the scMIC framework. (B) Enrichment of MICs in clone 1, which exhibits an aggressive phenotype. (C) Pseudotime (PseudoEMT) of primary tumor cells inferred by Monocle, showing a positive association with EMT. (D) Comparison of epithelial and mesenchymal gene expression between MICs and non-MICs. (E) Relationship between PseudoEMT and metastatic potential estimated by the scMIC framework.

Following this, pseudotime trajectory analysis of primary tumor cells inferred by Monocle revealed a strong correlation with EMT score, hereafter referred to as PseudoEMT (Figure 3C). This analysis demonstrated that MICs exhibited distinct EMT-associated features relative to non-MICs. Specifically, MICs showed reduced expression of epithelial markers and elevated expression of mesenchymal markers (Figure 3D, Figure S2C, Figure S3-4).

Notably, MIC-associated EMT patterns were similar across the data of two mouses. However, owing to differences in the number of reconstructed subclones and inter-individual heterogeneity, MICs in each mouse model also displayed distinct characteristics. For instance, the EMT-promoting gene S100A4 was markedly upregulated in MICs from mouse 2 (Figure S2C, Figure S4C), consistent with their aggressive clonal phenotype and in agreement with previous reports (6). Furthermore, closer examination of MICs in mouse 1 revealed that cells with the highest MIC scores were predominantly distributed in mid-to-late PseudoEMT stages, rather than in fully mesenchymal states (Figure 3E). This observation aligns with emerging evidence that metastatic potential peaks in rare, late hybrid EMT states.

Taken together, these analyses validated the reliability of scMIC in identifying MICs, and also highlighted its utility in uncovering key biological programs that drive metastatic initiation, such as hybrid EMT dynamics.

### 2.3 MICs relevant to clinical phenotypes

The identification of MICs is crucial for understanding their clinical relevance, particularly with regard to potential implications for metastasis diagnosis and treatment. To explore this, two scRNA-seq datasets (GSE178318 and OMIX002487) and one spatial transcriptomics dataset (GSE277783) were analyzed to investigate MICs or metastasis-initiating spots in primary tumors. In these human clinical datasets, tumor cells or spots were initially identified based on tumor markers, CNVs, and XGBoost malignancy scores (Figure S5–S7). Subsequently, scMIC was applied to classify these tumor cells or spots as MICs or non-MICs.

Analysis of these datasets revealed informative patterns in MIC abundance within primary tumors. Specifically, in the GSE178318 dataset, the MIC ratio in three colorectal cancer patients showed a significant correlation with the severity of liver metastasis, with higher MIC ratios associated with increased numbers and larger metastatic lesions (Figure 4A). Compared with tumor stage, primary tumor cell number, or lymph node metastasis status, the MIC ratio emerged as a more informative indicator of metastatic burden. This conclusion was further supported by an independent pancreatic cancer dataset (OMIX002487), in which higher MIC ratios were associated with larger liver metastases, irrespective of tumor stage (Figure 4B).

**Figure 4.**
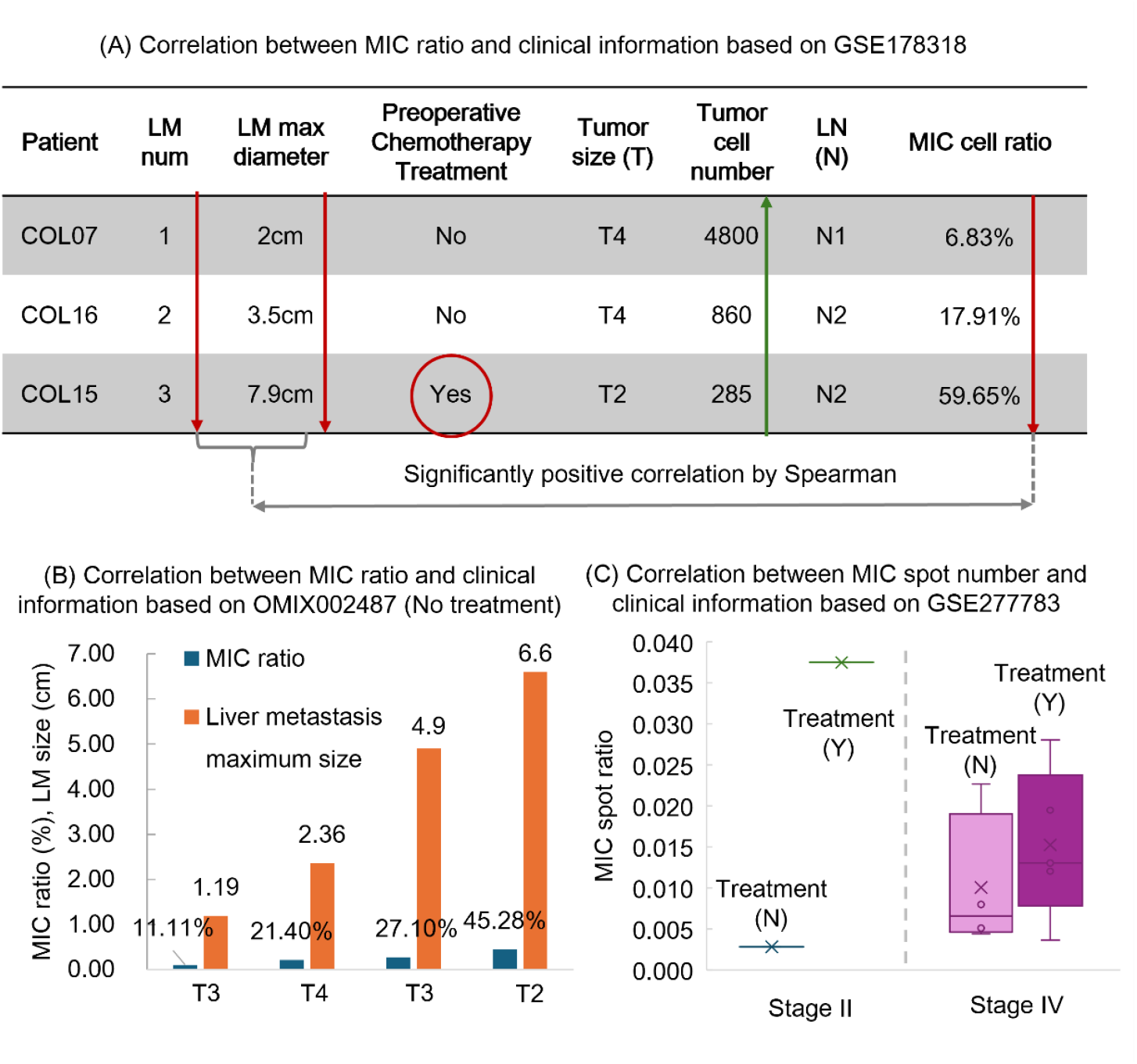
MICs associate with metastatic burden and treatment resistance. (A) The fraction of MICs positively correlates with both the size and number of liver metastases (LM) in the GSE178318 cohort. Notably, certain treatments are associated with a relative enrichment of MICs, as exemplified in COL15. (B) The MIC fraction shows a positive association with liver metastatic burden in the OMIX002487 cohort. (C) In the spatial transcriptomics dataset GSE277783, the proportion of MIC-positive spots increases following treatment. LN, lymph node.

Furthermore, the relationship between treatment and MIC abundance warrants attention. Specifically, an increased proportion of MICs was observed in primary tumors from colorectal cancer patients who received preoperative chemotherapy (Figure 4A). Consistently, spatial transcriptomic slides from treated pancreatic cancer patients also exhibited a higher proportion of MIC-associated spots (Figure 4C). This phenomenon may be partially explained by the differential impact of treatment on MICs and non-MICs. In particular, MICs exhibited significantly greater resilience than non-MICs across diverse microenvironmental contexts (19). Consequently, therapeutic interventions may preferentially eliminate non-MIC populations, leading to a relative enrichment of MICs within primary tumors. This shift could, in turn, increase the likelihood of metastatic dissemination. Collectively, these observations underscore the role of MICs in treatment resistance and highlight the importance of specifically targeting MICs in the development of effective therapeutic strategies.

Taken together, these analyses demonstrate the relevance of MIC abundance in primary tumors to clinical phenotypes, including metastatic burden and treatment response, across multiple cancer types and data modalities. In future clinical practice, MICs may offer new opportunities for metastasis risk stratification and therapeutic decision-making.

### 2.4 Gene programs underlying metastatic initiation

Given the strong associations between MICs and metastatic phenotype, as well as treatment efficacy, characterizing the intrinsic gene programs of MICs is of high biological relevance. To this end, we developed a framework combining an AutoEncoder with an MIC classifier (Figure 5A) to generate MIC-specific latent representations and identify associated gene programs. The framework was optimized through three stages, reconstruction loss, weighted classification loss, and merged loss (Figure 5A–B), to produce MIC representations for mouse 1 from the GSE173958 dataset (Figure 5C). Using the scRNA-seq expression matrix (Y) and the latent representations (L), a feature matrix (F) was derived to capture the associations between genes and latent variables. The training process was repeated ten times to ensure the robustness and reproducibility of MIC latent-associated genes. Finally, these genes were cross-referenced with genes highly expressed in MICs to define the final MIC gene sets.

**Figure 5.**
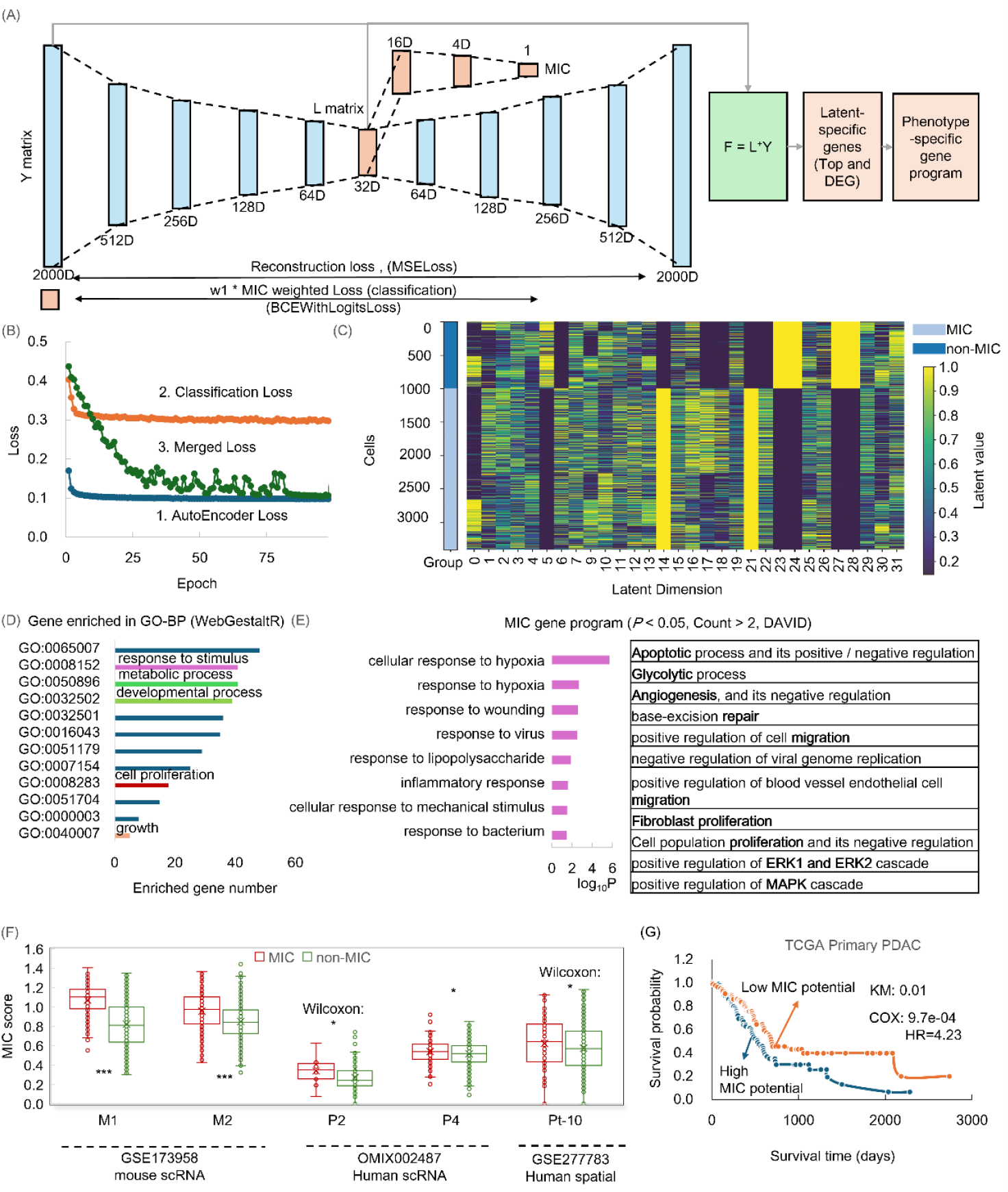
Pipeline for identifying the MIC signature. (A) Computational framework for identifying MIC latents and the MIC signature. (B) Three-stage training strategy for the computational framework. (C) Latent embeddings capturing MIC-related characteristics. (D) Genes contributing to MIC latents are enriched in biological processes such as response to stimulus. (E) Functional enrichment analysis using DAVID highlights involvement of these genes in specific programs, including response to diverse stimuli. (F) The MIC signature identified from M1 mouse data (GSE173958) is also highly enriched in additional mouse scRNA data (GSE173958), human scRNA data (OMIX002487), and human spatial transcriptomics data (GSE277783) tested by t-test or Wilcoxon test. (G) Patients with higher MIC signature scores exhibit poorer survival outcomes. ***, P < 0.001; **, P<0.05.

These MIC-associated genes were enriched in multiple biological pathways (Figure 5D–E). For instance, tumor cells rely on evasion of apoptosis to successfully establish metastases (20). Metabolic reprogramming, including enhanced glycolytic activity, represents a hallmark of metastatic progression and a potential therapeutic target (21). The ERK/MAPK signaling pathway plays a central role in driving cancer invasion and metastasis (22). Notably, responses to diverse stimuli were the most enriched pathways. These intrinsic alterations enhance the plasticity of MICs for fluctuating microenvironments adaptation and thereby facilitate metastatic dissemination. Collectively, the enrichment of these pathways underscores the fundamental characteristics of MICs.

To validate the utility of the MIC gene sets, we compared the average expression of these genes between MICs and non-MICs across multiple datasets, including scRNA-seq data from another mouse in GSE173958, human scRNA-seq data from OMIX002487, and human spatial transcriptomic spots from GSE277783. In most cases, MICs exhibited significantly higher average expression compared with non-MICs (Figure 5F). Furthermore, in bulk pancreatic cancer data from TCGA, patients with higher average expression of MIC-associated genes displayed poorer survival outcomes (Figure 5G). These findings highlight the gene program in driving metastatic initiation.

Given the demonstrated effectiveness of the MIC gene program, the approach was also applicable to primary tumor data when matched metastatic samples were unavailable. To evaluate this possibility, scRNA-seq data of mouse 2 from GSE173958 was analyzed. Cells with MIC program scores in the upper quartile were classified as MICs, whereas those in the lower quartile were classified as non-MICs (Figure S8A). Notably, 91.0% of MICs identified using this criterion overlapped with MICs defined based on paired primary–metastasis data. These cells exhibited the same aggressive clonal expansion and epithelial–mesenchymal characteristics (Figure S8B-C). This finding highlighted a strategy for identifying MICs directly from primary tumors.

### 2.5 Candidate biomarkers of MICs

Among the MIC-associated genes, OCIA domain–containing protein 2 (OCIAD2) has been reported to participate in responses to bacterial stimuli, maintain stem cell homeostasis, and regulate cell migration (23,24). Consistent with these established functions, the MICs identified in this study underscore a prominent role for OCIAD2 in metastatic progression (Figure 6A). Across mouse and human scRNA-seq datasets, as well as human spatial transcriptomic data, MICs exhibited higher OCIAD2 expression compared with non-MICs. Moreover, analysis of bulk tumor transcriptomic data revealed that OCIAD2 expression was significantly elevated in tumor tissues relative to normal controls (Figure 6B), was associated with poorer patient survival (Figure 6C), and correlated positively with cancer stemness (25) (Figure 6D). Collectively, these bioinformatic analyses highlight the potential involvement of OCIAD2 in driving metastasis initiating.

**Figure 6.**
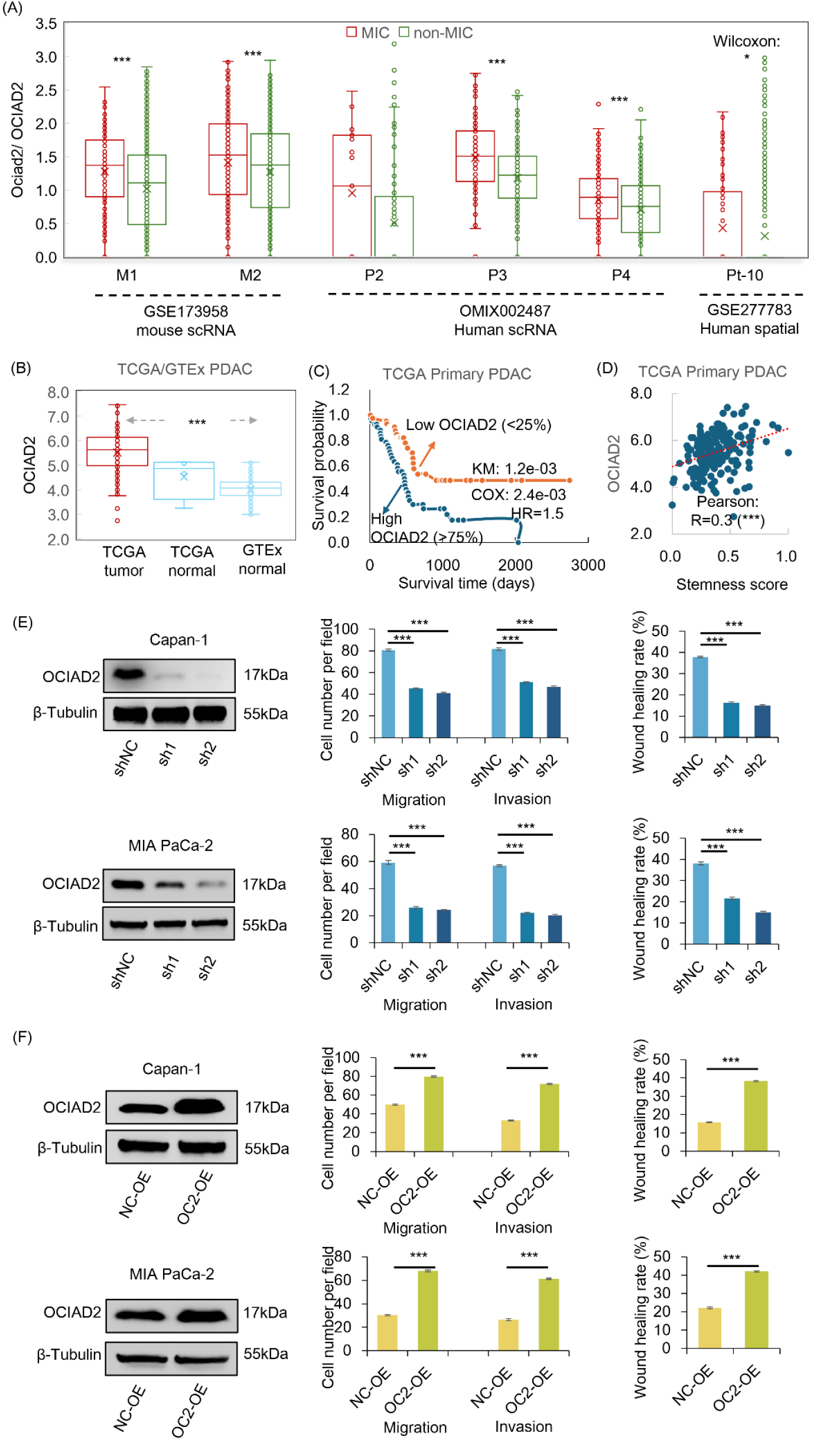
Effects of OCIAD2 on metastatic potential. (A) OCIAD2 is highly expressed in MICs for mouse scRNA-seq data (GSE173958), human scRNA-seq data (OMIX002487) and human spatial transcriptomics data (GSE277783) tested by t-test or Wilcoxon test. (B) OCIAD2 expression is elevated in bulk tumor tissues compared with normal tissues based on TCGA and GTEx datasets. (C) High OCIAD2 expression is associated with poorer overall survival. (D) OCIAD2 expression positively correlates with stemness scores. (E) Transwell migration and invasion assays, together with wound-healing assays, following OCIAD2 knockdown. (F) Transwell migration and invasion assays, together with wound-healing assays, following OCIAD2 overexpression.

To further elucidate the role of OCIAD2 in pancreatic cancer metastasis, we manipulated OCIAD2 expression in two pancreatic cancer cells, Capan-1 and MIA PaCa-2, using a lentivirus-mediated approach, employing shRNA for knockdown (shOCIAD2) and the pHBLV-UPP1 vector for overexpression (OCIAD2-OE). Cell migratory and invasive capacities were subsequently evaluated using transwell migration and invasion assays and wound healing assays. These assays revealed that OCIAD2 knockdown significantly impaired cell migration and invasion, whereas OCIAD2 overexpression markedly enhanced these metastatic phenotypes (Figure 6E–F, Figure S9–11). Collectively, these results demonstrate a pivotal role for OCIAD2 in promoting pancreatic cancer metastasis.

## 3. Discussion

Distinguishing MICs from the bulk of tumor cells is crucial for pinpointing the seeds of metastasis and elucidating underlying mechanisms. The scMIC framework identifies tumor cells in primary cancers with high metastatic potential by combining an embedding-based strategy, unbalanced optimal transport, and a top-k selection approach. To current knowledge, scMIC is the first algorithm to explicitly and reliably identify MICs. Existing methods infer MICs indirectly through pseudotime trajectories (7), stemness scores (8), pathway activities (9), or biomarker expression (10), often producing inconsistent results (Figure 2D–F) (6,11,12). This framework has been applied across multiple cancer types, including esophageal squamous cell carcinoma, colorectal cancer, and pancreatic cancer; across species, including mouse and human; and across various data types, including single-cell and spatial transcriptomics. These applications either demonstrate the reliability of the framework (Figures 2–3), propose potential clinical indicators of metastasis (Figure 4), or uncover previously uncharacterized mechanisms of metastasis (Figures 5–6). Collectively, these analyses highlight the utility of this framework for further metastasis research.

The successful identification of MICs in primary tumors (Figures 2–3) further reinforces the idea that the seeds of metastasis are already pre-established at the primary site (26). This finding provides a mechanistic foundation for the potential application of MIC-associated characteristics from primary tumors in clinical diagnosis and survival prognosis (Figures 4 and 5F – G). A particularly notable feature is the pronounced transcriptomic alterations in stimulus-response pathways (Figures 5D – E), reflecting enhanced cellular plasticity. This plasticity enables dynamic adaptation to environmental stresses and facilitates the acquisition of metastatic-initiating capacity (27). This inherent characteristic of MICs ultimately contributes to drug resistance and refractoriness to metastasis (28). Furthermore, growing evidence suggests treatment-induced metastasis (29), where one potential underlying mechanism is that treatment may increase the frequency of cells exhibiting metastatic phenotypes, such as those in MICs (Figure 4). Taken together, these observations underscore the critical role of MIC research in advancing metastasis diagnosis and therapeutic strategies.

The scMIC framework is primarily evaluated and applied using paired primary and metastatic tumor samples from the same individuals (Figure 2 – 6). To broaden its applicability in scenarios where only primary tumor data are available, two complementary strategies are proposed. First, the MIC gene program derived from paired primary and metastatic samples can be directly applied to other primary tumor datasets lacking metastatic samples, as shown in Figure S8A. Second, for each query primary tumor sample, the nearest reference neighbor cells can be identified using scFoundation (30) embeddings combined with a KNN search (Figure S12A–B). These reference neighbors carry MIC or non-MIC labels assigned by scMIC on paired datasets, enabling accurate label transfer to the query cells. Both strategies were validated using the primary tumor data from mouse 2 in the GSE173958 dataset and produced results consistent with those obtained from paired datasets (Figure S8B–C, Figure S12C–D).

The migration process is shaped by both the tumor “seed” and the microenvironment of the secondary organ. This study examines the shared transcriptomic dynamics of tumor cells during metastasis, enabling the identification of cells or patients with higher metastatic potential (Figure 4–6). Such insights are valuable for assessing metastatic risk and guiding clinical decision-making. Additionally, the metastatic “soil” also plays criticalm role in supporting successful colonization. Further investigation is needed to determine which secondary organs individual cells or patients are most predisposed to metastasize to. Developing a foundation model that integrates bulk, single-cell, and spatial transcriptomics across primary tumors and multiple metastatic sites may offer a promising approach to address this question.

## 4. Materials and methods

### 4.1 Primary and metastatic data

The MIC identification and characterization framework was developed and evaluated using multiple cancer metastasis datasets. Model training was performed on scRNA-seq data from a multi-timepoint esophageal squamous cell carcinoma mouse model (GSE249057). Further validation was conducted using scRNA-seq data from a metastatic pancreatic cancer mouse model with annotated clonal aggressiveness (GSE173958). Clinical relevance was assessed using human colorectal cancer scRNA-seq data (GSE178318) (27), human pancreatic cancer scRNA-seq data (OMIX002487), and human pancreatic cancer spatial transcriptomics data (GSE277783) (28) with detailed clinical annotations such as metastasis number, metastasis size, and treatment status. All the pancreatic cancer–related datasets were further integrated with TCGA bulk tumor data, and in vitro cell line experiments to characterize MIC-associated gene programs and to explore candidate MIC biomarkers.

### 4.2 Data preprocessing

All scRNA-seq datasets were pre-processed following a standardized workflow, including quality control, data normalization, dimensionality reduction, cell clustering, tumor cell identification, and differential expression analysis across tumor subclusters.

For the GSE249057 scRNA-seq dataset, cells were filtered based on quality control criteria with more than 200 and fewer than 8,000 detected genes, more than 1,000 total RNA counts, and a mitochondrial gene fraction below 15%. Gene expression counts were then scaled to 10,000 per cell and log-transformed to generate the normalized expression matrix. Subsequently, PCA was performed for dimensionality reduction, and the top 50 PCs were used for cell clustering with the Louvain algorithm. As this dataset consisted exclusively of GFP-positive, flow-sorted tumor cells, explicit tumor cell identification was not required. Finally, differentially expressed genes (DEGs) across tumor subclusters were identified using FindAllMarkers, and genes with P < 0.05 and log^2^ fold change > 0 were retained for downstream MIC identification.

For the GSE173958 scRNA-seq dataset, which contains fluorescence-sorted tumor cells, pre-processing followed the same workflow as described above. Given that the deposited dataset comprised high-quality cells expressing more than 1,000 genes, quality control filtering was limited to retaining cells with mitochondrial gene fractions below 15%. Subsequently, normalization, dimensionality reduction, clustering, and differential expression analysis were performed using the same parameters as described above.

For the GSE178318 scRNA-seq datasets, pre-processing followed the same general workflow with dataset-specific modifications. Following quality control filtering, cells with 500–20,000 UMI counts, more than 200 detected genes, and mitochondrial gene fractions below 15% were retained. Subsequently, normalization, dimensionality reduction, and clustering were performed as described above. Tumor cells were then identified based on the expression of epithelial markers EPCAM, KRT8, KRT18, and KRT19, in combination with CNV-based and XGBoost-derived malignancy scores implemented in scCancer2 (18). Finally, DEGs were identified across tumor subclusters for downstream analyses.

For the OMIX002487 scRNA-seq dataset, only annotated epithelial cells were retained for downstream analysis. Following normalization and clustering, tumor cells were identified based on the expression of EPCAM, KRT8, KRT18, and KRT19 together with inferred CNV patterns computed using CopyKAT (31), consistent with the original study. Subsequently, DEGs were identified across tumor subclusters.

For the GSE277783 spatial transcriptomics dataset, spots with UMI counts below 100 were excluded. Following this, normalization, dimensionality reduction, and clustering were performed using the same approaches applied to the scRNA-seq datasets. Tumor-cell-enriched spots were then identified by integrating inferred CNV scores computed using SPATA2 (32), exceeding those observed in normal spots, with enrichment of tumor lineage – specific transcriptional markers. To further improve robustness, CNV-based and XGBoost-based malignancy scores were additionally applied. Finally, DEGs were identified across tumor subclusters.

### 4.3 Metastasis-initiating cells identification framework

A subset of primary tumor cells may undergo intrinsic reprogramming, enabling them to better adapt to diverse microenvironments during metastasis and successfully colonize secondary organs. These cells, referred to as MICs, are hypothesized to exhibit stronger associations with metastatic tumor cells than other primary tumor cells (13). To investigate this process, UOT (16) was applied to infer the transport plan between primary and metastatic tumor cells (Figure 1A).

Within the framework, expression-based associations were jointly leveraged to characterize cellular relationships. The transport plan *γ*\* ∈ ℝ ^*n×m*^ was obtained by minimizing the following objective:

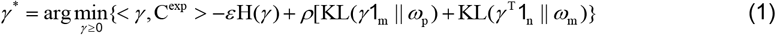

Where 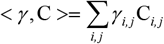 denotes the Frobenius inner product representing the total transport cost, C^e×p^ ∈ ℝ^*n*×*m*^ is the gene expression–based cost matrix defined as the Euclidean distance between PCA or AutoEncoder representations of primary and metastatic tumor expression profile, 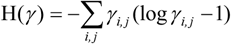 is the entropy term with regularization coefficient ϵ>0, KL(· ‖ ·) denotes the Kullback – Leibler divergence controlling relaxation of row- (*ω*_p_) and column-wise (*ω*_m_) marginals with regularization coefficient ρ>0. The resulting optimal γ thus defines the transport plan, mapping each metastatic tumor cell (columns) to its corresponding primary tumor cells (rows).

Since each metastatic cell is assumed to originate from a single primary tumor cell, a top-k filtering scheme with k=1 was applied to γ to enhance both interpretability and sparsity. Specifically, only the largest transport weight in each column of γ, corresponding to each metastatic cell, was retained. The metastatic potential of each primary tumor cell (row) was then calculated by summing the weights across the corresponding columns of γ, alternatively unnormalized or normalized by either the total or the maximum value across all primary tumor cells.

### 4.4 Gene program identification framework for Metastasis-initiating cells

To investigate the molecular mechanisms underlying metastasis initiation, a framework combining an AutoEncoder with a multilayer perceptron (MLP) MIC classifier was developed to extract MIC-specific latent representations and gene programs (Figure 5A). The AutoEncoder took the scRNA-seq matrix of highly variable genes as both input and output, learning low-dimensional latent embeddings that capture key transcriptional patterns. These embeddings were then fed into the MLP-based MIC classifier, which predicted MIC labels previously identified by scMIC. By combining unsupervised latent learning with supervised MIC classification, this framework maintains the interpretability of the latent embeddings.

The framework was trained on scRNA-seq data from mouse 1 (GSE173958) using a three-stage optimization procedure repeated ten times. The stages sequentially minimized the reconstruction loss, the MIC classifier loss, and a combined loss. The overall loss function is defined as:

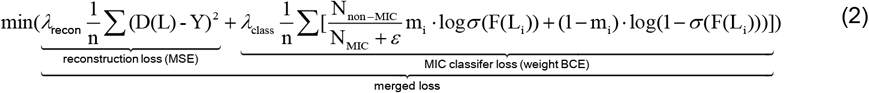

where λ balances the contributions of each component, D(·) is the decoder network, L represents the latent embedding, Y is the scRNA-seq expression matrix, N is the number of MIC or non-MIC cells, ϵ=1×10^−6^, m denotes the MIC or non-MIC label, σ(·) is the sigmoid activation function, and F represents the MLP-based MIC classifier.

Using the learned latent embeddings together with the original scRNA-seq data, the feature matrix F was computed as:

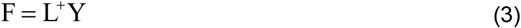

where L^+^ denotes the Moore-Penrose pseudoinverse of the latent matrix L. This feature matrix captures associations between genes and latent representations. From this matrix, genes were selected for the MIC-specific program if they ranked among the top 500 contributors to the MIC latents in all ten training repetitions and were highly expressed in MICs (adjusted P < 0.05, log_2_ fold change > 0.1, percentage > 0.5). Functional characterization of this gene program was performed using WebGestaltR and the DAVID enrichment tool. An average expression score of these genes was calculated as the “MIC potential,” which was then validated across other pancreatic cancer datasets included in this study.

### 4.5 Biomarker discover for Metastasis-initiating cells

Within the MIC-specific gene program, Ociad2 was selected for further investigation of its role in metastasis initiation. Its expression, along with that of its human homolog OCIAD2, was assessed across MICs and non-MICs in multiple pancreatic cancer datasets. In addition, its expression was evaluated in bulk tumor samples relative to normal controls, and its association with patient survival outcomes and tumor stemness was examined. Collectively, these analyses provide systematic evidence supporting the relevance of OCIAD2 to metastasis initiation across multi-scale datasets. Finally, in vitro cell line experiments were conducted to further validate its functional role in metastasis, as described in detail below.

The human pancreatic cancer cell lines MIA PaCa-2 and Capan-1 were obtained from the State Key Laboratory of Holistic Integrative Management of Gastrointestinal Cancers at Xijing Hospital. MIA PaCa-2 cells were cultured in DMEM (Gibco) supplemented with 10% fetal bovine serum (FBS, Gibco), while Capan-1 cells were cultured in IMDM medium (Gibco) supplemented with 20% FBS (Gibco). Stable cell lines were generated via lentiviral transduction to express shOCIAD2-NC, shOCIAD2-KD1, and shOCIAD2-KD2, as well as pHBLV-con (designated as OCIAD2-NC) and pHBLV-OCIAD2 (referred to as OCIAD2-OE). All lentiviral constructs were synthesized by Genechem (Shanghai, China).

Cells were lysed in RIPA lysis buffer (NCMBiotech, WB3100) supplemented with protease and phosphatase inhibitors (NCMBiotech, P002) and incubated on ice for 15 min. Lysates were centrifuged at 12,000 rpm for 15 min at 4 ° C, and the protein-containing supernatants were collected. Equal amounts of protein were mixed with SDS–PAGE sample loading buffer (Beyotime, P0015) and denatured by boiling at 100 °C for 10 min.

Proteins were separated on YoungPAGE gels (GenScript, M00930) using MOPS running buffer and subsequently transferred onto PVDF membranes (Millipore, IPVH00010) using a wet transfer system. Membranes were blocked with PVA to prevent non-specific binding and incubated overnight at 4 °C with primary antibodies. After washing, membranes were incubated with the corresponding secondary antibodies for 2 h at room temperature.

Protein bands were visualized using the e-BLOT imaging system. Primary antibodies used included anti-human OCIAD2 (Proteintech, 13437-1-AP, 1:2000) and anti-β-tubulin (Proteintech, HRP-60008, 1:5000). The secondary antibody used was anti-rabbit IgG (Cell Signaling Technology, 7074S, 1:5000).

Wound healing and Transwell assays were performed to assess cell migratory capacity. For the Transwell migration assay, cells were seeded into 24-well Transwell chambers (Corning, NY, USA) at a density of 5 × 10^4^ cells per well in 200 μL serum-free medium. After incubation, migrated cells were fixed with 4% paraformaldehyde solution and stained with 0.25% crystal violet. For the wound healing assay, transfected pancreatic cancer cells were cultured in six-well plates until approximately 80% density. A linear wound was created by vertically scratching using a 200 μL pipette, followed by replacement with 2 mL serum-free medium. Wound closure was monitored and quantified by measuring the average wound width over time.

## 5. Data availability

All data used in this study were obtained from the GEO (GSE249057, GSE173958, GSE178318, and GSE277783), NGDC (OMIX002487), TCGA, and GTEx databases.

## 6. Code availability

All code associated with this study is available at https://github.com/swu13/scMIC.

## 7. Supplementary Data

Supplementary files are available online along with the manuscript.

## 8. Acknowledgement

This work was supported by the National Natural Science Foundation of China (Grant No. 82227802, 62002270) and the Fundamental Research Funds for the Central Universities.

## References

1. Gerstberger, S., Jiang, Q. and Ganesh, K. (2023) Metastasis. Cell, 186, 1564–1579.

2. Lambert, A.W., Zhang, Y. and Weinberg, R.A. (2024) Cell-intrinsic and microenvironmental determinants of metastatic colonization. Nat. Cell Biol., 26, 687–697.

3. Fares, J., Fares, M.Y., Khachfe, H.H., Salhab, H.A. and Fares, Y. (2020) Molecular principles of metastasis: a hallmark of cancer revisited. Signal Transduct Target Ther, 5, 28.

4. Karras, P., Black, J.R., McGranahan, N. and Marine, J.-C. (2024) Decoding the interplay between genetic and non-genetic drivers of metastasis. Nature, 629, 543–554.

5. Wong, C.N., Zhang, Y., Ru, B., Wang, S., Zhou, H., Lin, J., Lyu, Y., Qin, Y., Jiang, P., Lee, V.H. et al. (2024) Identification and Characterization of Metastasis-Initiating Cells in ESCC in a Multi-Timepoint Pulmonary Metastasis Mouse Model. Advanced science (Weinheim, Baden-Wurttemberg, Germany), 11, e2401590.

6. Simeonov, K.P., Byrns, C.N., Clark, M.L., Norgard, R.J., Martin, B., Stanger, B.Z., Shendure, J., McKenna, and Lengner, C.J. (2021) Single-cell lineage tracing of metastatic cancer reveals selection of hybrid EMT states. Cancer Cell, 39, 1150–1162 e9.

7. Wang, G., Shi, C., He, L., Li, Y., Song, W., Chen, Z., Liu, Z., Wang, Y., He, X., Yu, Y. et al. (2024) Identification of the tumor metastasis-related tumor subgroups overexpressed NENF in triple-negative breast cancer by single-cell transcriptomics. Cancer Cell Int., 24, 319.

8. Li, R., Liu, X., Huang, X., Zhang, D., Chen, Z., Zhang, J., Bai, R., Zhang, S., Zhao, H., Xu, Z. et al. (2024) Single-cell transcriptomic analysis deciphers heterogenous cancer stem-like cells in colorectal cancer and their organ-specific metastasis. Gut, 73, 470–484.

9. Yang, C., Li, Y., Wang, Z., Shan, H., Zhang, G., Meng, X., Wang, G., Hou, Z., Zhao, X., Zhang, X. et al. (2025) Identification of a cancer stem cell-like subpopulation that promotes HCC metastasis. JHEP reports : innovation in hepatology, 7, 101302.

10. Hiraga, T., Ito, S. and Nakamura, H. (2013) Cancer stem-like cell marker CD44 promotes bone metastases by enhancing tumorigenicity, cell motility, and hyaluronan production. Cancer Res., 73, 4112–22.

11. Fico, F., Bousquenaud, M., Rüegg, C. and Santamaria-MartÍnez, A. (2019) Breast Cancer Stem Cells with Tumor-versus Metastasis-Initiating Capacities Are Modulated by TGFBR1 Inhibition. Stem cell reports, 13, 1–9.

12. Pham, D.X. and Hsu, T. (2025) Tumor-initiating and metastasis-initiating cells of clear-cell renal cell carcinoma. J. Biomed. Sci., 32, 17.

13. Li, G., Béal, E., Sumner, D., Galli, G.G., Cremasco, V., Korn, J.M., Dondelinger, F., Ruddy, D., Kauffmann, and Dimitrieva, S. (2024) Predicting metastatic transcriptomes of patient tumors with deep learning. Cancer Res., 84, 896–896.

14. Cang, Z., Zhao, Y., Almet, A.A., Stabell, A., Ramos, R., Plikus, M.V., Atwood, S.X. and Nie, Q. (2023) Screening cell–cell communication in spatial transcriptomics via collective optimal transport. Nat. Methods, 20, 218–228.

15. Bunne, C., Schiebinger, G., Krause, A., Regev, A. and Cuturi, M. (2024) Optimal transport for single-cell and spatial omics. Nature Reviews Methods Primers, 4, 58.

16. Cang, Z. and Nie, Q. (2020) Inferring spatial and signaling relationships between cells from single cell transcriptomic data. Nat Commun, 11, 2084.

17. Kolouri, S., Park, S.R., Thorpe, M., Slepcev, D. and Rohde, G.K. (2017) Optimal mass transport: Signal processing and machine-learning applications. IEEE signal processing magazine, 34, 43–59.

18. Chen, Z., Miao, Y., Tan, Z., Hu, Q., Wu, Y., Li, X., Guo, W. and Gu, J. (2024) scCancer2: data-driven in-depth annotations of the tumor microenvironment at single-level resolution. Bioinformatics, 40.

19. Massagué, J. and Ganesh, K. (2021) Metastasis-Initiating Cells and Ecosystems. Cancer Discov., 11, 971–994.

20. Su, Z., Yang, Z., Xu, Y., Chen, Y. and Yu, Q. (2015) Apoptosis, autophagy, necroptosis, and cancer metastasis. Mol. Cancer, 14, 48.

21. Wei, Q., Qian, Y., Yu, J. and Wong, C.C. (2020) Metabolic rewiring in the promotion of cancer metastasis: mechanisms and therapeutic implications. Oncogene, 39, 6139–6156.

22. Bahar, M.E., Kim, H.J. and Kim, D.R. (2023) Targeting the RAS/RAF/MAPK pathway for cancer therapy: from mechanism to clinical studies. Signal Transduct Target Ther, 8, 455.

23. Yin, Y.F., Jia, Q.Y., Yao, H.F., Zhu, Y.H., Zheng, J.H., Duan, Z.H., Hu, C.Y., Sun, Y.W., Liu, D.J., Huo, Y.M. et al. (2024) OCIAD2 promotes pancreatic cancer progression through the AKT signaling pathway. Gene, 927, 148735.

24. Consortium, A.o.G.R. (2020) Alliance of Genome Resources Portal: unified model organism research platform. Nucleic Acids Res., 48, D650–D658.

25. Zhou, W., Su, M., Jiang, T., Xie, Y., Shi, J., Ma, Y., Xu, K., Xu, G., Li, Y. and Xu, J. (2024) Cancer Stemness Online: A Resource for Investigating Cancer Stemness and Associations with Immune Response. Genomics Proteomics Bioinformatics, 22.

26. Sanghvi, N., Calvo-Alcañiz, C., Rajagopal, P.S., Scalera, S., Canu, V., Sinha, S., Schischlik, F., Wang, K., Madan, S., Shulman, E. et al. (2024) Charting the transcriptomic landscape of primary and metastatic cancers in relation to their origin and target normal tissues. Sci. Adv., 10, eadn0220.

27. Pérez-González, A., Bévant, K. and Blanpain, C. (2023) Cancer cell plasticity during tumor progression, metastasis and response to therapy. Nature cancer, 4, 1063–1082.

28. Shi, X., Wang, X., Yao, W., Shi, D., Shao, X., Lu, Z., Chai, Y., Song, J., Tang, W. and Wang, X. (2024) Mechanism insights and therapeutic intervention of tumor metastasis: latest developments and perspectives. Signal Transduct Target Ther, 9, 192.

29. Ebos, J.M. (2015) Prodding the Beast: Assessing the Impact of Treatment-Induced Metastasis. Cancer Res., 75, 3427–35.

30. Hao, M., Gong, J., Zeng, X., Liu, C., Guo, Y., Cheng, X., Wang, T., Ma, J., Zhang, X. and Song, L. (2024) Large-scale foundation model on single-cell transcriptomics. Nat. Methods, 21, 1481–1491.

31. Gao, R., Bai, S., Henderson, Y.C., Lin, Y., Schalck, A., Yan, Y., Kumar, T., Hu, M., Sei, E., Davis, A. et al. (2021) Delineating copy number and clonal substructure in human tumors from single-cell transcriptomes. Nat. Biotechnol., 39, 599–608.

32. Kueckelhaus, J., Frerich, S., Kada-Benotmane, J., Koupourtidou, C., Ninkovic, J., Dichgans, M., Beck, J., Schnell, O. and Heiland, D.H. (2024) Inferring histology-associated gene expression gradients in spatial transcriptomic studies. Nat Commun, 15, 7280.

