## Supplementary Figures1-13 for "Identification and Characterization of Metastasis-initiating cells"

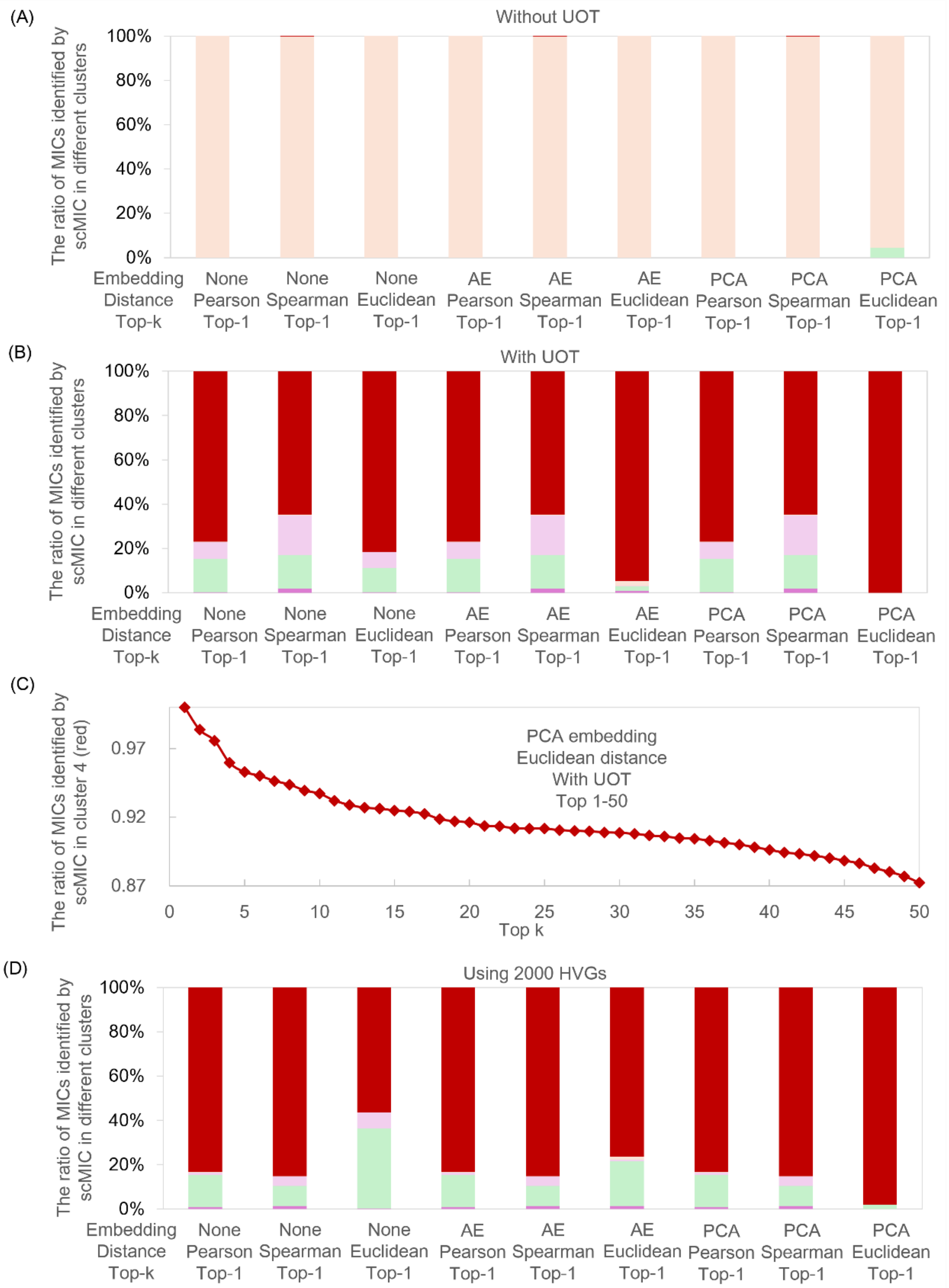


Figure S1. Evaluation of the scMIC framework under different analytical settings. (A) Performance of scMIC without unbalanced optimal transport (UOT). (B) Performance of scMIC across different embedding strategies and distance metrics. (C) Performance of scMIC under varying Top-k selection thresholds. (D) Performance of scMIC using the top 2,000 highly variable genes (HVGs). All other analyses in this study were performed using differentially expressed genes from primary tumor subclusters. PCA, principal component analysis; AE, autoencoder.

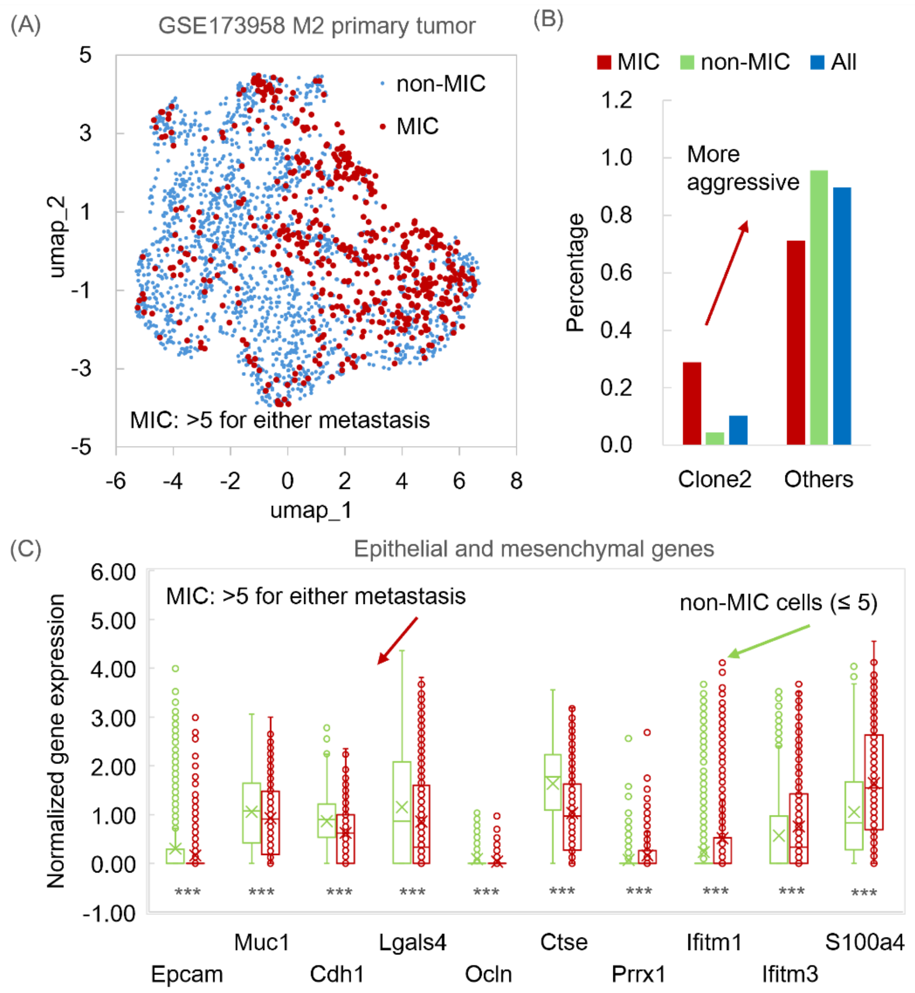


Figure S2. MICs associate with an aggressive clone in a lineage-traced M2 mouse model from the GSE173958 dataset. (A) UMAP visualization of primary tumor cells with MICs identified by the scMIC framework. (B) Enrichment of MICs in clone 2, which exhibits an aggressive phenotype. (C) Comparison of epithelial and mesenchymal gene expression between MICs and non-MICs.

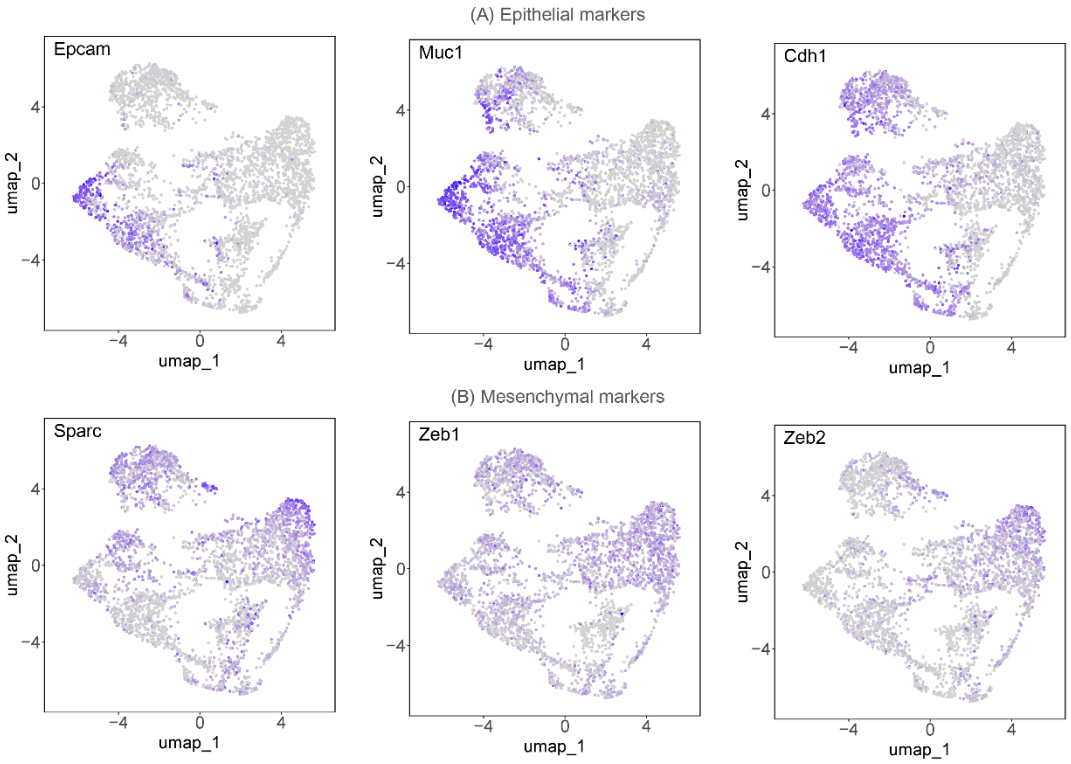


Figure S3. UMAP visualization of EMT marker expression in a lineage-traced M1 mouse model from the GSE173958 dataset. (A) UMAP visualization of epithelial marker expression. (B) UMAP visualization of mesenchymal marker expression.

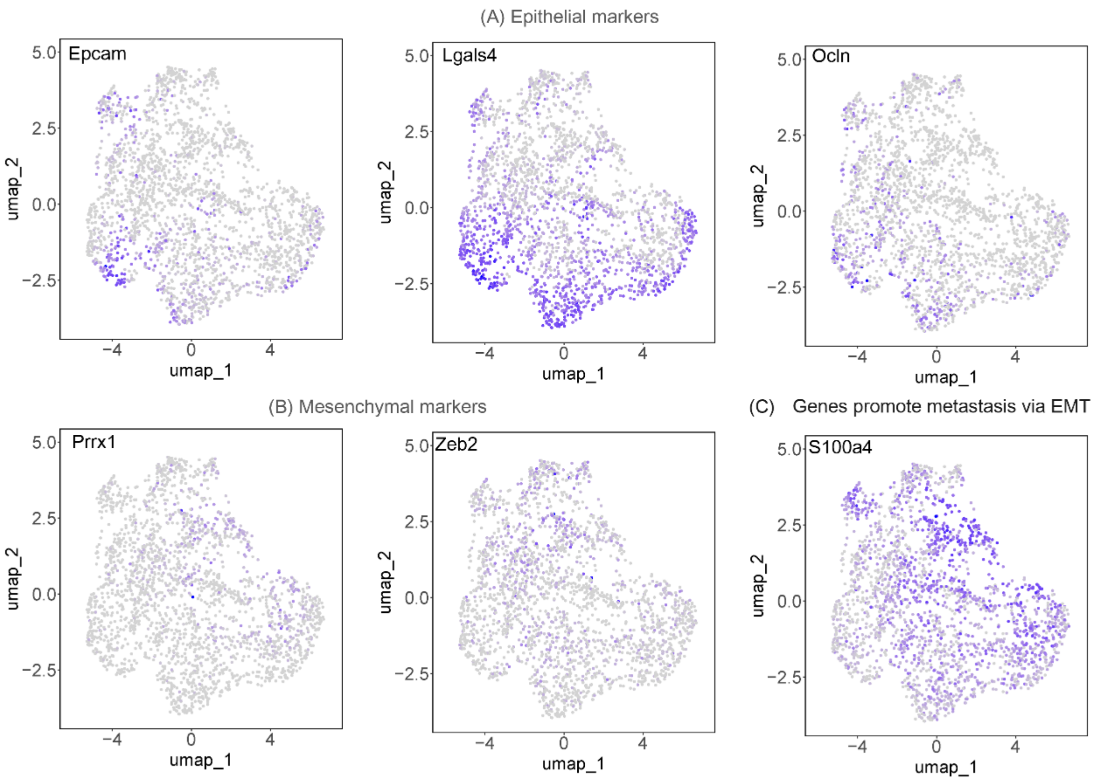


Figure S4. UMAP visualization of EMT and metastasis-related marker expression in a lineage-traced M2 mouse model from the GSE173958 dataset. (A) UMAP visualization of epithelial marker expression. (B) UMAP visualization of mesenchymal marker expression. (C) UMAP visualization of a metastasis-related marker expression.

(A)

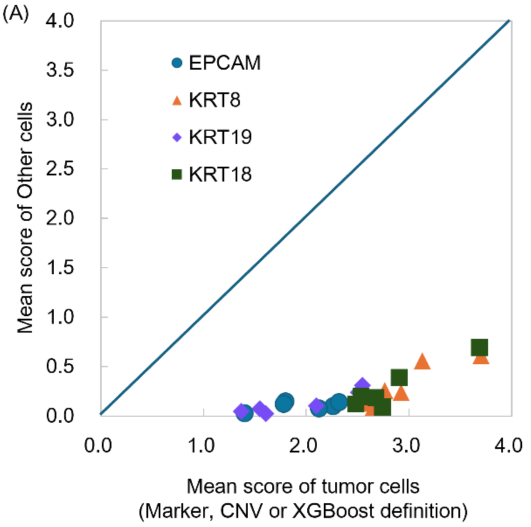


(B)

| Patient | Sample | Tumor cells number |
|---|---|---|
| COL07 | Primary | 4,800 |
| | Metastasis | 939 |
| COL15 | Primary | 285 |
| | Metastasis | 491 |
| COL16 | Primary | 860 |
| | Metastasis | 207 |

Figure S5. Identification of tumor cells in the GSE178318 dataset. (A) Expression of canonical epithelial markers (EPCAM, KRT8, KRT18, and KRT19) in tumor cells compared with other cell populations, as identified by marker-based, CNV-based, or XGBoost (scCancer) approaches. (B) Number of tumor cells identified in each sample.

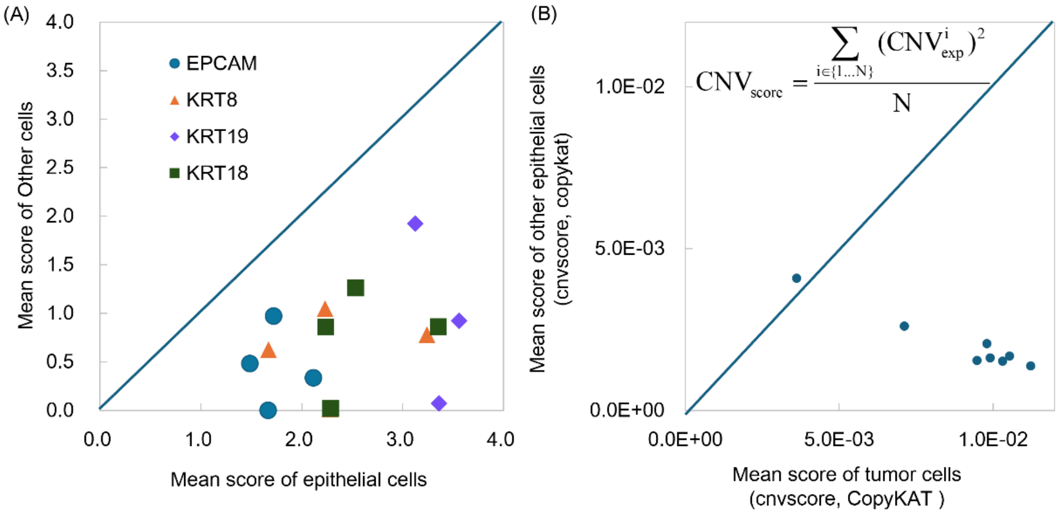


(C)

| Group | Tumor cells number | | | |
|---|---|---|---|---|
| | P2 | P3 | P4 | P5 |
| Primary | 135 | 417 | 860 | 8,281 |
| Metastasis | 175 | 169 | 2,443 | 10,441 |

Figure S6. Identification of tumor cells in the OMIX002487 dataset. (A) Expression of canonical markers (EPCAM, KRT8, KRT18, and KRT19) in epithelial cells compared with other cell populations. (B) Average CNV scores in tumor cells identified using the CopyKAT approach compared with other epithelial cells enriched for canonical markers. (B) Number of tumor cells identified in each sample.

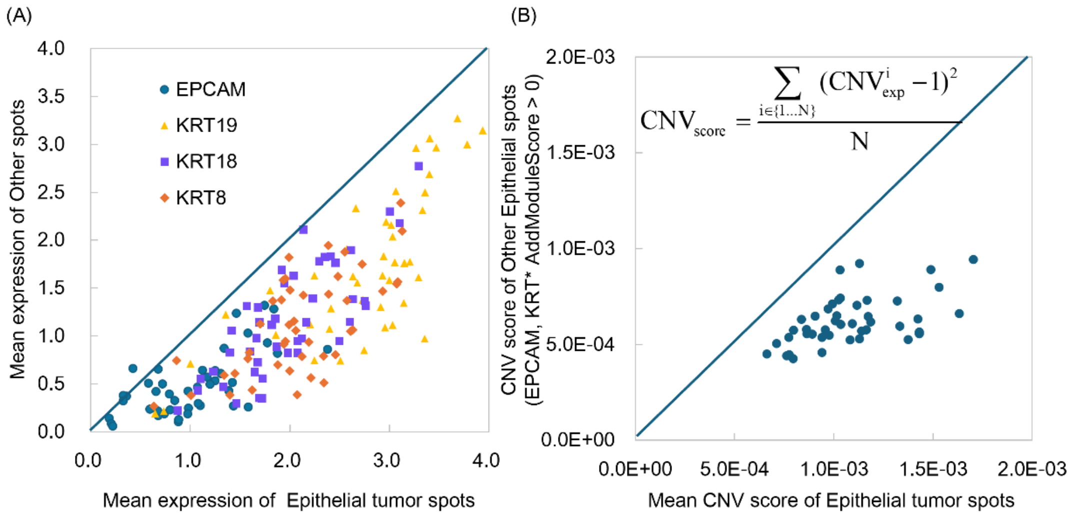


1. EPCAM,KRT8,KRT19,KRT18 Geneset AddModuleScore > 0
2. SPATA2: InferCNV score > reference CNV score
3. scCancer: malignant based on CNV or XGBoost identification

(C)

| Sample | TumorSpot | Sample | TumorSpot | Sample | TumorSpot | Sample | TumorSpot | Sample | TumorSpot |
|---|---|---|---|---|---|---|---|---|---|
| Pt-1A | 1,429 | Pt-4B | 2,715 | Pt-6D | 1,615 | Pt-9A | 2,326 | Pt-11B | 1,026 |
| Pt-1C | 2,117 | Pt-4C | 2,492 | Pt-7A | 1,642 | Pt-9B | 2,647 | Pt-11C | 1,146 |
| Pt-2A | 661 | Pt-4D | 3,410 | Pt-7B | 3,168 | Pt-9C | 2,027 | Pt-11D | 404 |
| Pt-2B | 1,903 | Pt-5A | 2,720 | Pt-7C | 1,755 | Pt-9D | 1,378 | Pt-13A | 2,067 |
| Pt-2C | 395 | Pt-5B | 2,339 | Pt-7D | 2,726 | Pt-10A | 962 | Pt-13B | 174 |
| Pt-3A | 2,937 | Pt-5C | 2,363 | Pt-8A | 3,606 | Pt-10B | 1,381 | Pt-13C | 578 |
| Pt-3B | 1,753 | Pt-6A | 1,532 | Pt-8B | 2,448 | Pt-10C | 1,165 | Pt-13D | 617 |
| Pt-3C | 1,786 | Pt-6B | 813 | Pt-8C | 2,673 | Pt-10D | 1,478 | Pt-13E | 1,147 |
| Pt-4A | 2,884 | Pt-6C | 1,577 | Pt-8D | 159 | Pt-11A | 400 | | |

Figure S7. Identification of tumor spots in the GSE277783 dataset. (A) Expression of canonical epithelial markers (EPCAM, KRT8, KRT18, and KRT19) in tumor spots compared with other spots, as identified by marker-based, CNV-based (SPATA2 and scCancer), or XGBoost (scCancer) approaches. (B) Average CNV scores in tumor spots compared with other epithelial spots showing enriched canonical markers. (B) Number of tumor spots identified in each sample.

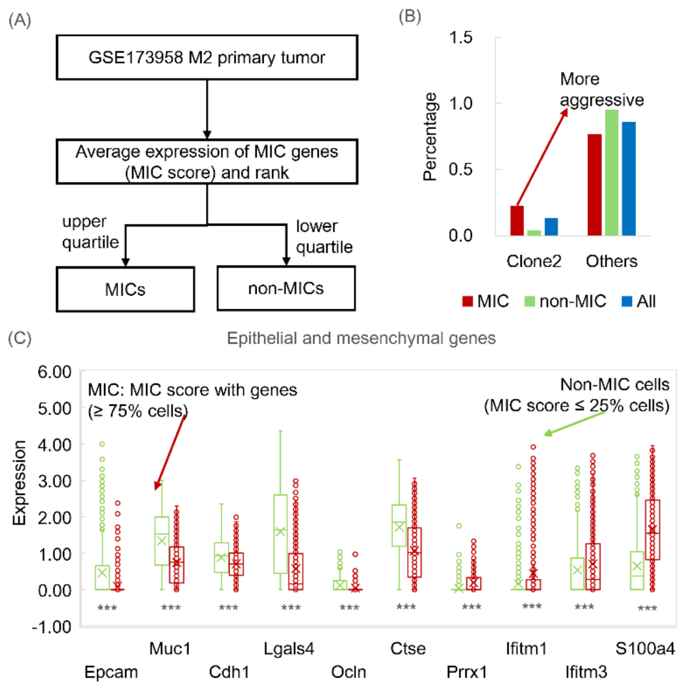


Figure S8. Effectiveness of MIC identification using MIC programs. (A) Definition of MICs according to MIC programs. (B) Distribution of clones between MICs and non-MICs as determined by MIC programs. (C) Comparison of epithelial and mesenchymal gene expression between MICs and non-MICs identified by MIC programs.

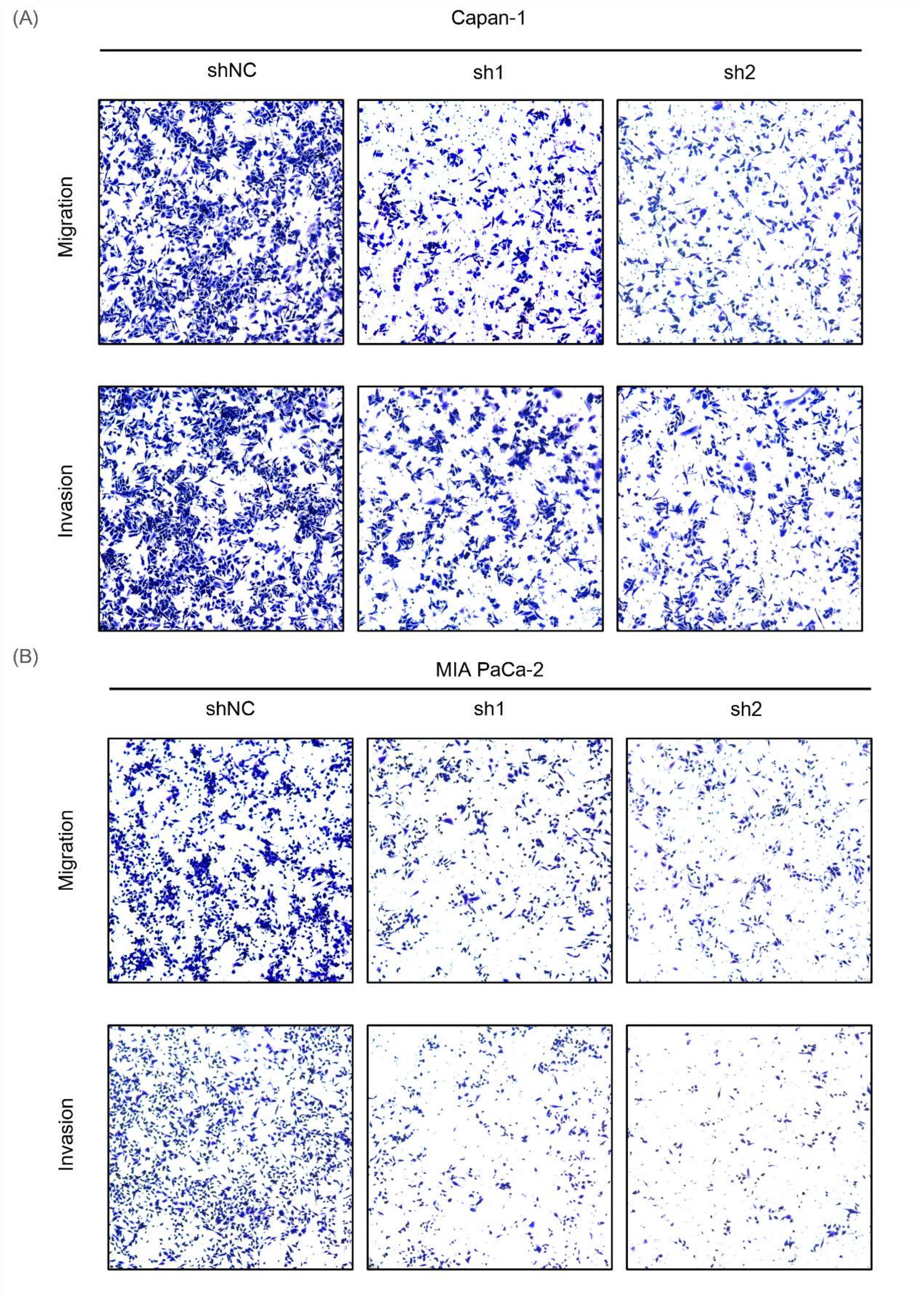


Figure S9. Transwell migration and invasion assay results of Capan-1 (A) and MIA PaCa-2 (B) cells following lentivirus-mediated OCIAD2 knockdown using shRNA (shOCIAD2).

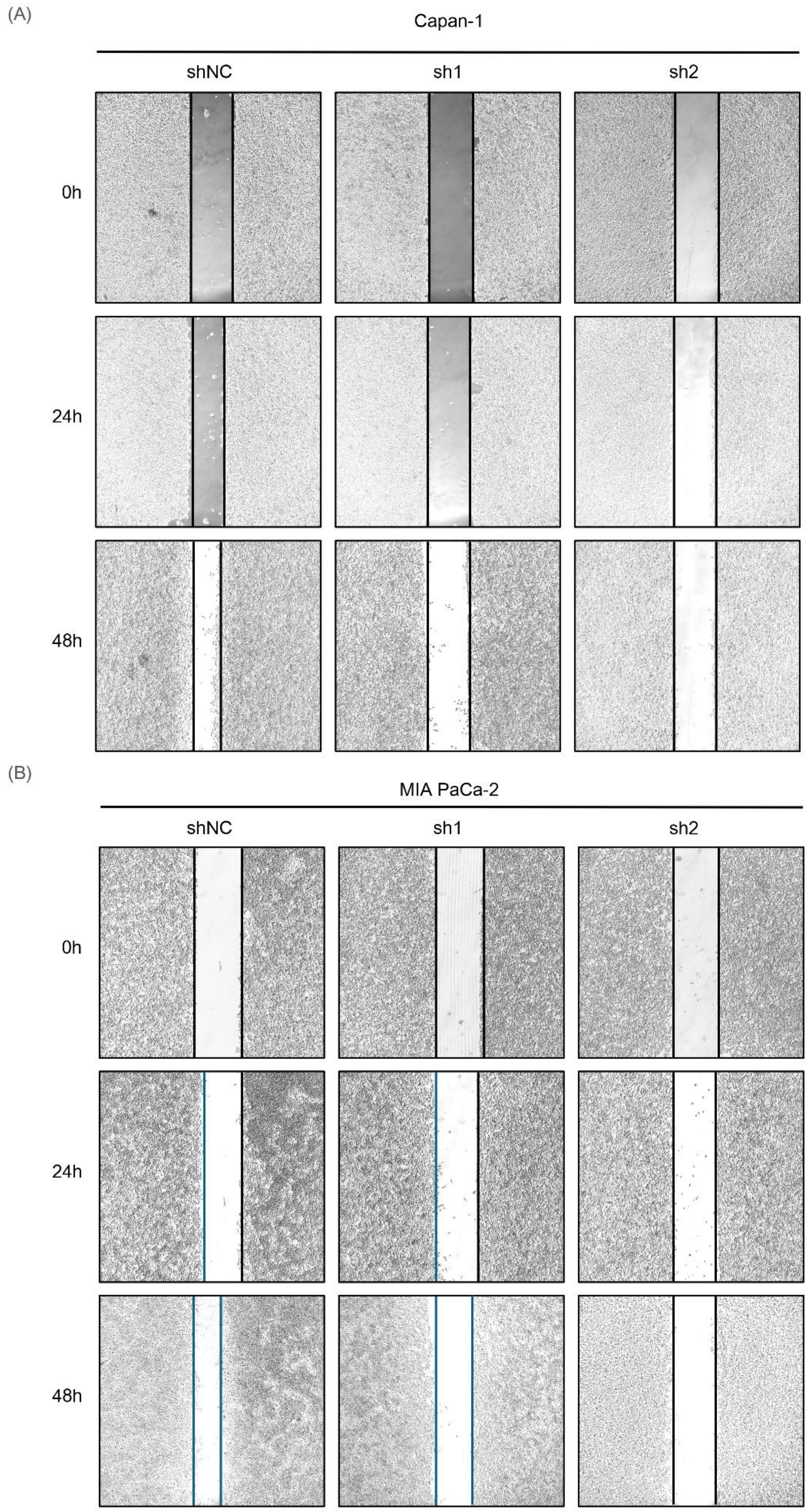


Figure S10. Wound healing assay results of Capan-1 (A) and MIA PaCa-2 (B) cells following lentivirus-mediated OCIAD2 knockdown using shRNA (shOCIAD2).

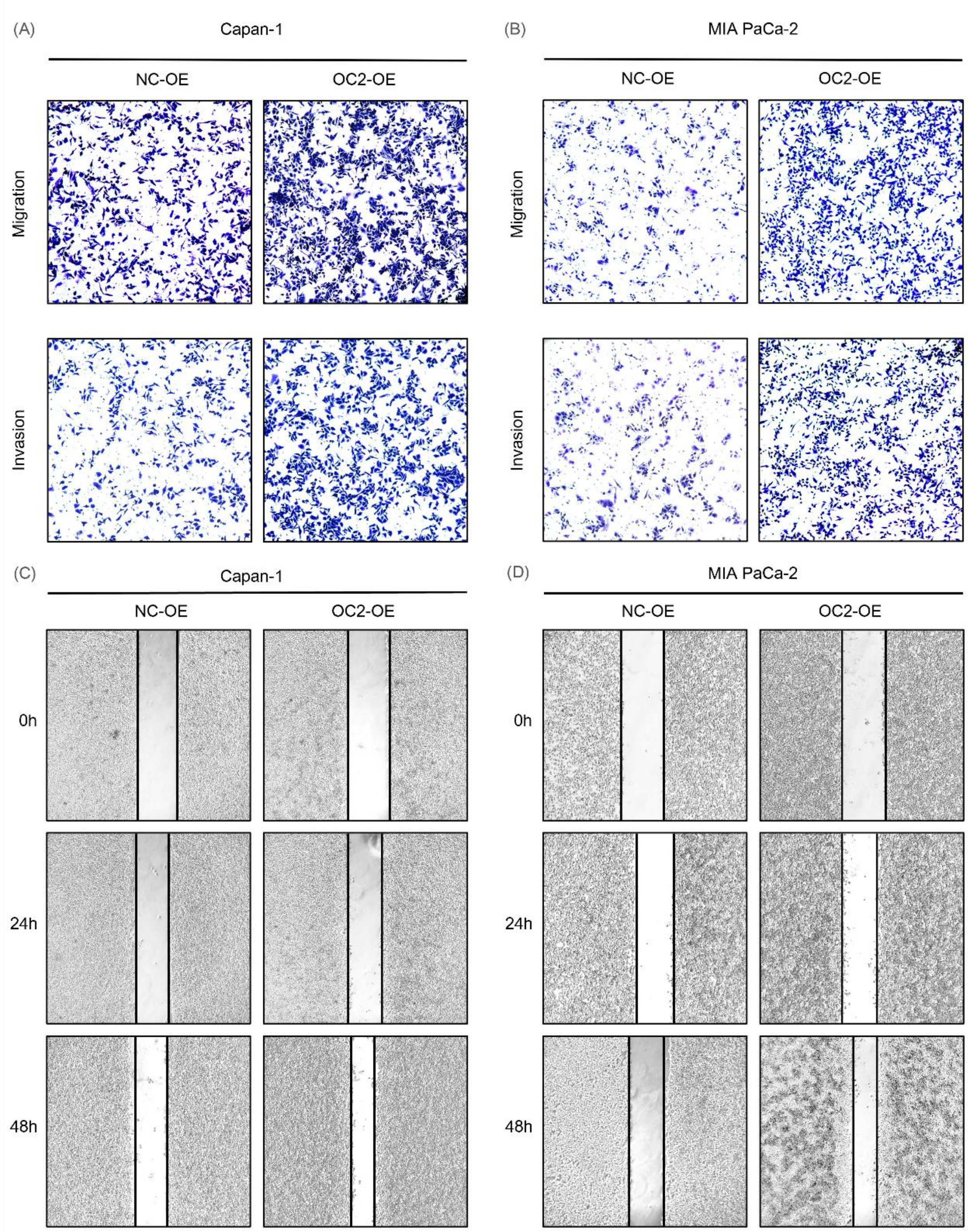


Figure S11. Effects of OCIAD2 overexpression on PDAC cell migration and invasion. Transwell migration and invasion assay results of Capan-1 (A) and MIA PaCa-2 (B) cells following OCIAD2 overexpression (OCIAD2-OE). Wound healing assay results of Capan-1 (C) and MIA PaCa-2 (D) cells following OCIAD2 overexpression (OCIAD2-OE).

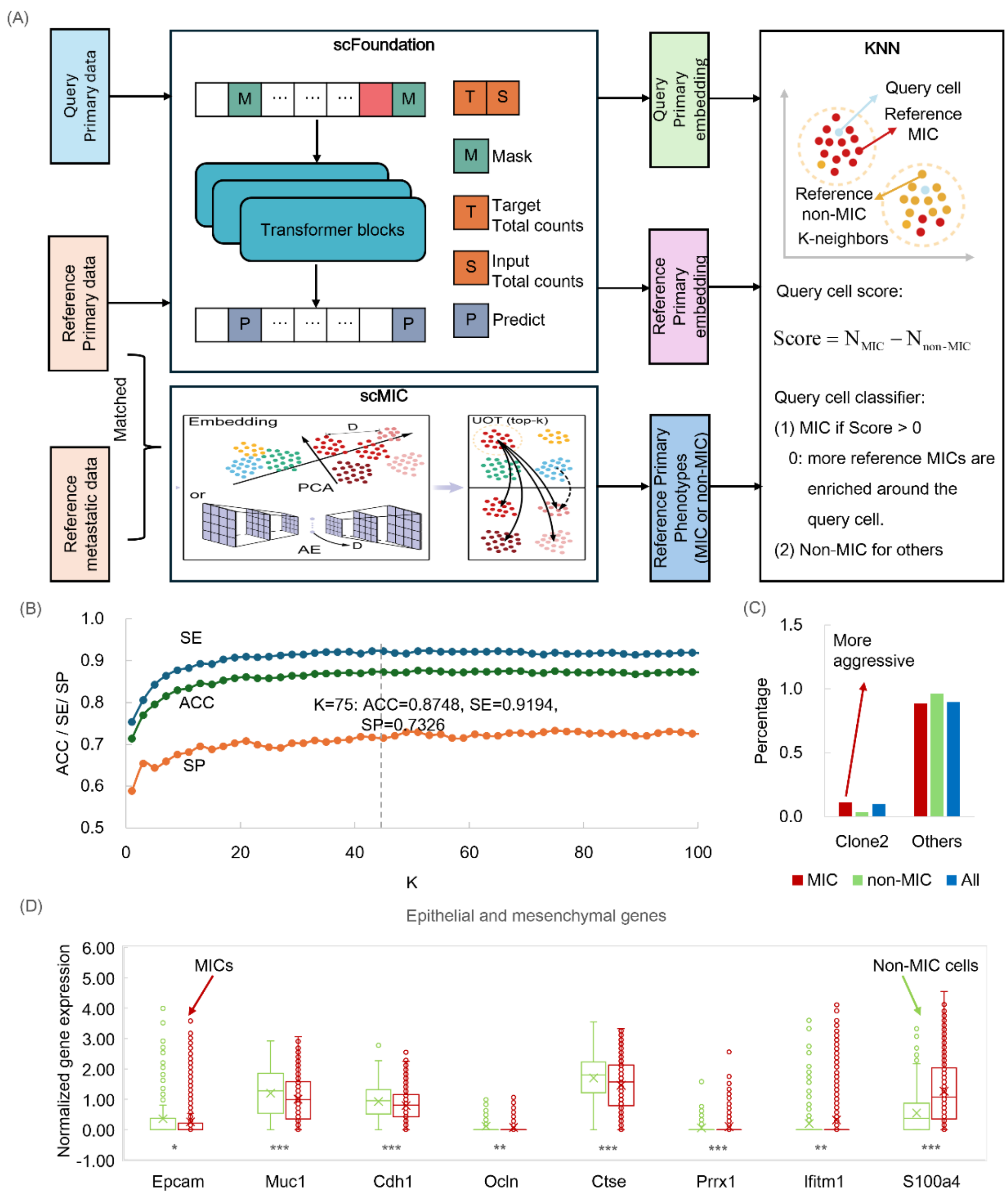


Figure S12. Identification of MICs using primary data only. (A) Workflow for identifying MICs in query primary samples by integrating matched primary–metastasis reference data with scFoundation and K-Nearest Neighbors (KNN). (B) Performance of the model evaluated using the primary samples from the M2 mouse in the GSE173958 dataset. MICs and non-MICs defined by scMIC using matched primary–metastasis data serve as the gold standard. (C) Clonal distribution of predicted MICs and non-MICs based solely on primary data. (D) Comparison of epithelial and mesenchymal gene expression between predicted MICs and non-MICs using primary data only. SE, sensitivity; SP, specificity; ACC, accuracy.
